# Mechanism of FtsZ assembly dynamics revealed by filament structures in different nucleotide states

**DOI:** 10.1101/2021.11.29.470395

**Authors:** Federico M. Ruiz, Sonia Huecas, Alicia Santos-Aledo, Elena A. Prim, José M. Andreu, Carlos Fernández-Tornero

**Affiliations:** Centro de Investigaciones Biológicas Margarita Salas CSIC, Ramiro de Maeztu 9, 28040 Madrid, Spain

## Abstract

Treadmilling protein filaments perform essential cellular functions by growing from one end while shrinking from the other, driven by nucleotide hydrolysis. Bacterial cell division relies on the primitive tubulin homolog FtsZ, a target for antibiotic discovery that assembles into single treadmilling filaments that hydrolyse GTP at an active site formed upon subunit association. We determined high-resolution filament structures of FtsZ from the pathogen *Staphylococcus aureus* in complex with different nucleotide analogues and cations, including mimetics of the ground and transition states of catalysis. Together with mutational and biochemical analyses, our structures reveal interactions made by the GTP γ-phosphate and Mg^2+^ at the subunit interface, a K^+^ ion stabilizing loop T7 for co-catalysis, new roles of key residues at the active site and a nearby crosstalk area, and rearrangements of a dynamic water shell bridging adjacent subunits upon GTP hydrolysis. We propose a mechanistic model that integrates nucleotide hydrolysis signalling with assembly-associated conformational changes and filament treadmilling. Equivalent assembly mechanisms may apply to more complex tubulin and actin cytomotive filaments that share analogous features with FtsZ.

## Introduction

FtsZ is an assembling GTPase that plays a key role during bacterial cell division [1]. This widely-conserved protein polymerizes in the presence of GTP and metal cations into polar filaments that gather at the centre of dividing cells to form a dynamic Z-ring [2]. FtsZ filaments associate with a variable set of partner proteins into the divisome, which coordinates membrane invagination and cell wall peptidoglycan synthesis during cytokinesis [3,4]. As such, bacterial cell division proteins are targets for discovering new antibiotics [5].

The head-to-tail treadmilling dynamics of FtsZ filaments, observed both *in vitro* [6,7] and *in vivo* [8–10], depends on two major properties of this protein (see scheme in Fig 1A). First, GTP hydrolysis only occurs within the filament, as the complete catalytic site is formed at the interface between adjacent FtsZ monomers [11,12]. Second, assembly of the FtsZ filament relies on the existence of two conformations that are independent of the bound nucleotide [13,14]. The relaxed (R) conformation present in free monomers exhibits low affinity for the filament, while the tense (T) conformation is acquired by filament subunits. Switching between these two conformations explains cooperative assembly of single stranded filaments [15,16] and creates different pairs of encountering interfaces at each filament end, thus enabling kinetic polarity for treadmilling [14,17]. GTP-bound FtsZ monomers associate to the filament growing end and, once inside the filament, are retained in the T conformation due to simultaneous contact with subunits on both of its sides, even after GTP hydrolysis. The subunit at the shrinking end dissociates due to weakened contact with its only neighbouring GDP-bound subunit (Fig 1A). FtsZ filaments shrink from the exposed nucleotide end, as deduced from mutational studies [17,18]. A recent numerical model generates populations of treadmilling filaments that accurately reproduce experimental results, thus unifying the described FtsZ properties [19].

**Fig 1.**
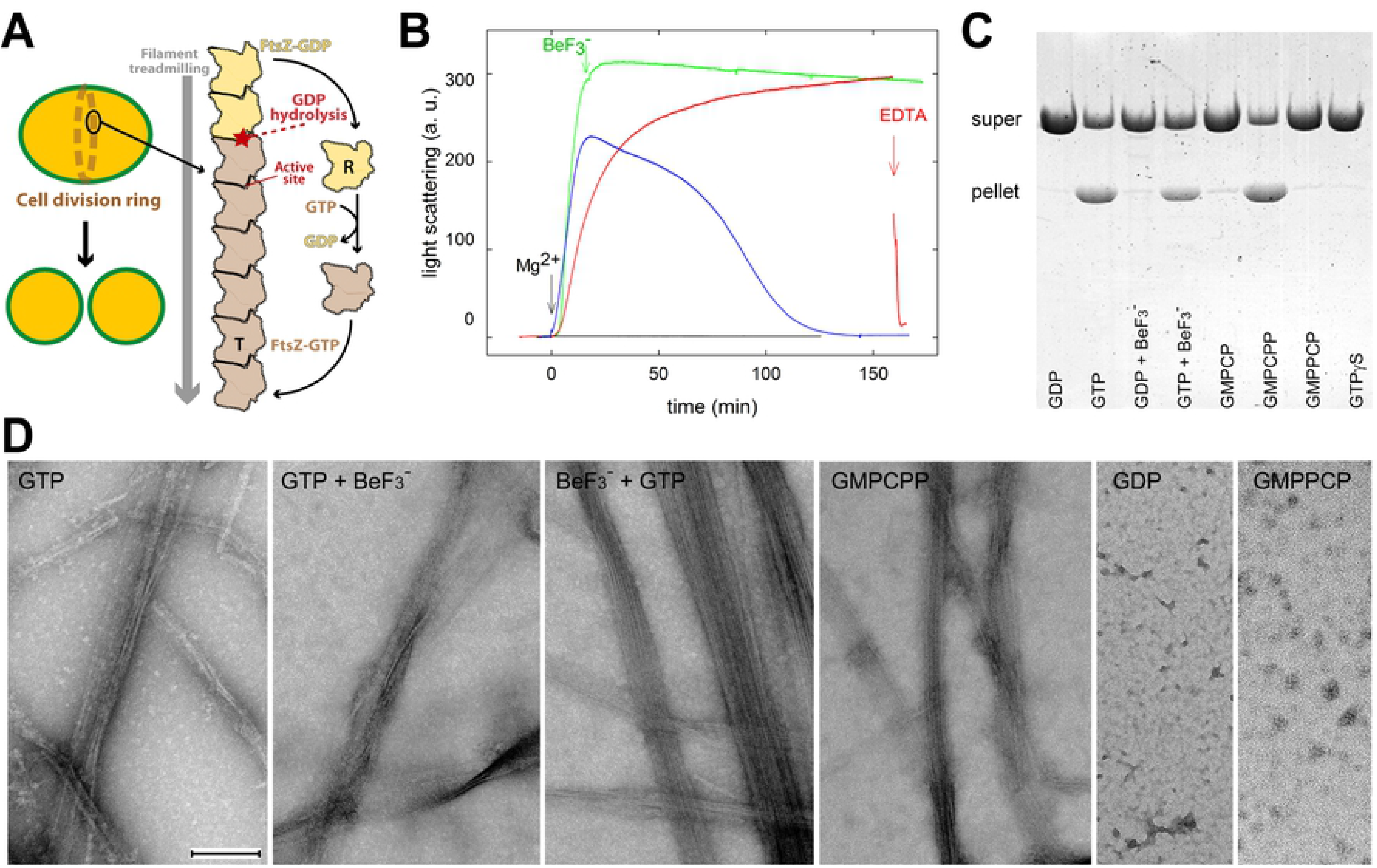
SaFtsZ filament assembly with GTP mimetics. (A) Schematic representation of the Z-ring, made of FtsZ filament clusters, in dividing *S. aureus* cells, next to a treadmilling FtsZ filament, drawn with only eight subunits for simplicity. Brown and yellow subunits are bound to GTP and GDP, respectively, while their conformational state is indicated by “T” for the filament and “R” for the monomer. (B) Stabilization of SaFtsZ assembly by BeF3- in solution. Light scattering at 350 nm was employed to monitor the formation of polymers by the SaFtsZ core (50 µM) with 1 mM GTP (blue line), 1 mM GTP plus 1 mM BeF_3_^-^ (red line), 1 mM GTP to which 1 mM BeF_3_^-^ was added at the point indicated by the arrow (green line; named GTP+BeF_3_^-^ below), and 1 mM GDP plus 1 mM BeF_3_^-^ (black line). Assembly was triggered by addition of 10 mM MgCl_2_ at time zero. Samples containing both GTP and 1 mM BeF_3_^-^ disassembled with 20 mM EDTA (indicated by the red arrow). (C) Sedimentation of polymers formed by SaFtsZ (31 µM) with different guanine nucleotides and mimetics (1 mM each, except 0.1 mM GMPCP and GMPCPP); pellet and supernatant samples were consecutively loaded with a 15 min time shift in SDS-PAGE. (D) Representative electron micrographs of polymers formed by SaFtsZ (31 µM) with different nucleotide mimetics, in negative stain. Incubation time was 30 min with GTP and GTP+BeF_3_^-^ (green line in panel A), 24 h with BeF_3_^-^+GTP added 5 min later, 80 min with GMPCPP, 2 h with GDP and GMPPCP, to visualize polymer structures at maximal formation. The bar indicates 200 nm.

FtsZ consists of an enzymatic core followed by a flexible C-terminal tail that contains binding motifs for membrane-tethering proteins [3]. The primitive enzymatic core of FtsZ, conserved with tubulin [20], comprises an N-terminal nucleotide-binding domain (NBD) and a C-terminal GTPase-activating domain (GAD) connected by the long central H7 *α*-helix [12,21]. The nucleotide occupies a pocket in the NBD, where loops T1 to T4 contact the phosphate groups, loop T5 interacts with the ribose moiety, and loop T6 contributes to guanine binding. At the opposite face of the monomer, the T7 synergy loop contains two conserved catalytic aspartates that promote GTP hydrolysis of the nucleotide placed in the adjacent subunit within the filament [11,12]. While loop T7 exhibits remarkable plasticity in FtsZ structures in the R conformation [13,14,22–24], it adopts a defined arrangement by coordinating a metal ion in the T conformation [25]. Concomitant rotation of the neighbouring GAD enables the formation of straight filaments where the nucleotide is buried from the solvent. The antibacterial compound PC190723 and chemically-related FtsZ inhibitors bind in a cleft between NBD and GAD of subunits in the T conformation, thus blocking filament disassembly [25–27]. A reduced interaction area in the R conformation leads to monomers [14] or pseudofilaments where the nucleotide is partly exposed to the solvent [13,23,24]. Crystal structures of FtsZ filaments in the T conformation are only available in the presence of GDP, GTP with a truncated non-catalytic T7 loop, or the non-hydrolysable GTP analogue GTPγS [13,25,28,29]. Therefore, mechanistic information on GTP hydrolysis and its implication in filament assembly dynamics is currently largely missing.

We report a dozen filament structures of the FtsZ core from the pathogen *S. aureus*, hereafter SaFtsZ, in complex with various GTP mimetics and metal ions. The high resolution of these structures enables description of key interactions at the interface of adjacent monomers forming the catalytic site with unprecedented detail. Together with structure-based mutational analysis of critical residues around the active site, our results shed light on the mechanisms of FtsZ filament assembly, GTP hydrolysis and treadmilling.

## Results

### Stabilization of FtsZ polymers by GTP mimetics and cations

We analysed the effect of several GTP mimetics on SaFtsZ (residues 12-316) assembly, monitoring polymer formation with light scattering and sedimentation assays (Fig 1B and 1C). SaFtsZ assembly required GTP plus MgCl_2_ and polymers disassembled upon nucleotide exhaustion. BeF_3_^-^ is a chemical mimetic of phosphate that acts on G-proteins [30,31] and tubulin [32,33] as a reversibly-binding analogue of the GTP *γ*-phosphate. We found that addition of BeF_3_^-^ stabilizes SaFtsZ polymers against disassembly, suggesting that BeF_3_^-^ replaces the *γ*-phosphate in SaFtsZ filaments following GTP hydrolysis. Fast depolymerization can be induced by Mg^2+^ chelation with EDTA. BeF_3_^-^ with GDP and Mg^2+^ was ineffective for polymer assembly. Negative-staining electron microscopy showed that similar bundles of SaFtsZ filaments (Fig 1D) form with GTP, GTP plus BeF_3_^-^ and GMPCPP, a slowly-hydrolysing analogue of GTP that induces robust SaFtsZ assembly [29]. The non-hydrolysable GTP analogues GMPPCP and GTP*γ*S fail to induce SaFtsZ assembly in these experiments, similarly to GDP and its analogue GMPCP, highlighting their inability to mimic GTP function on SaFtsZ (Fig 1C and 1D). We confirmed that BeF_3_^-^ stabilizes the assembly of full-length SaFtsZ in a qualitatively similar manner to the SaFtsZ core, as well as that of distant homologs from *Escherichia coli* (EcFtsZ) and *Methanocaldococcus jannaschii* (MjFtsZ) (S1 Fig). Repolymerization with BeF_3_^-^ after disassembly was only observed for MjFtsZ, possibly related to residual GTP.

Aluminium fluoride (AlF_x_) is a mimetic of the transition state of GTP catalysis, as described for classical GTPases [31,34]. We observed that, in contrast to BeF_3_^-^, AlF_3_/AlF_4_^-^ (AlF_x_) with GDP and Mg^2+^ induces SaFtsZ to form short, C-shaped, single and double filaments of ∼100 nm in diameter (S2 Fig). These distinct AlF_x_ polymers form rapidly with GDP and more slowly with GTP, possibly reflecting the need of GTP hydrolysis for AlF_x_ binding in place of the γ-phosphate. The striking filament curvature [35], which is similar to the ∼150 nm circular filaments observed during FtsZ disassembly [36], likely relates to structural alterations at the association interface between SaFtsZ monomers in solution.

### Structure determination of FtsZ filaments with GTP mimetics

Our attempts to determine the structure of SaFtsZ with bound GTP in the T conformation employing X-ray crystallography yielded filament structures where density in the active site only accounted for GDP, like in reported structures [25,29]. Therefore, we solved twelve crystal structures of SaFtsZ in complex with different GTP mimetics, analogues and metal ions, at resolutions ranging from 1.4 to 1.9 Å (S1 Table), enabling unambiguous assignment of ligand densities (S3 Fig). We complemented this study with four additional structures of single-residue mutants complexed to GDP (S1 Table). In all cases, SaFtsZ adopts the T conformation forming straight filaments with minor but relevant differences between them, as detailed below. Subsequent description assumes a filament orientation where nucleotide-binding loops T1-T6 in the NBD face upwards, while the T7 synergy loop in the GAD looks downwards (Fig 2A). Four major contact regions can be defined in longitudinal contacts between top and bottom monomers (Fig 2B): (A) top helix H10 with bottom loop H6-H7; (B) top strand S9 with bottom loop T5; (C) top helix H8 with bottom loop T3; (D) top loop T7 with bottom helices H1 and H2.

**Fig 2.**
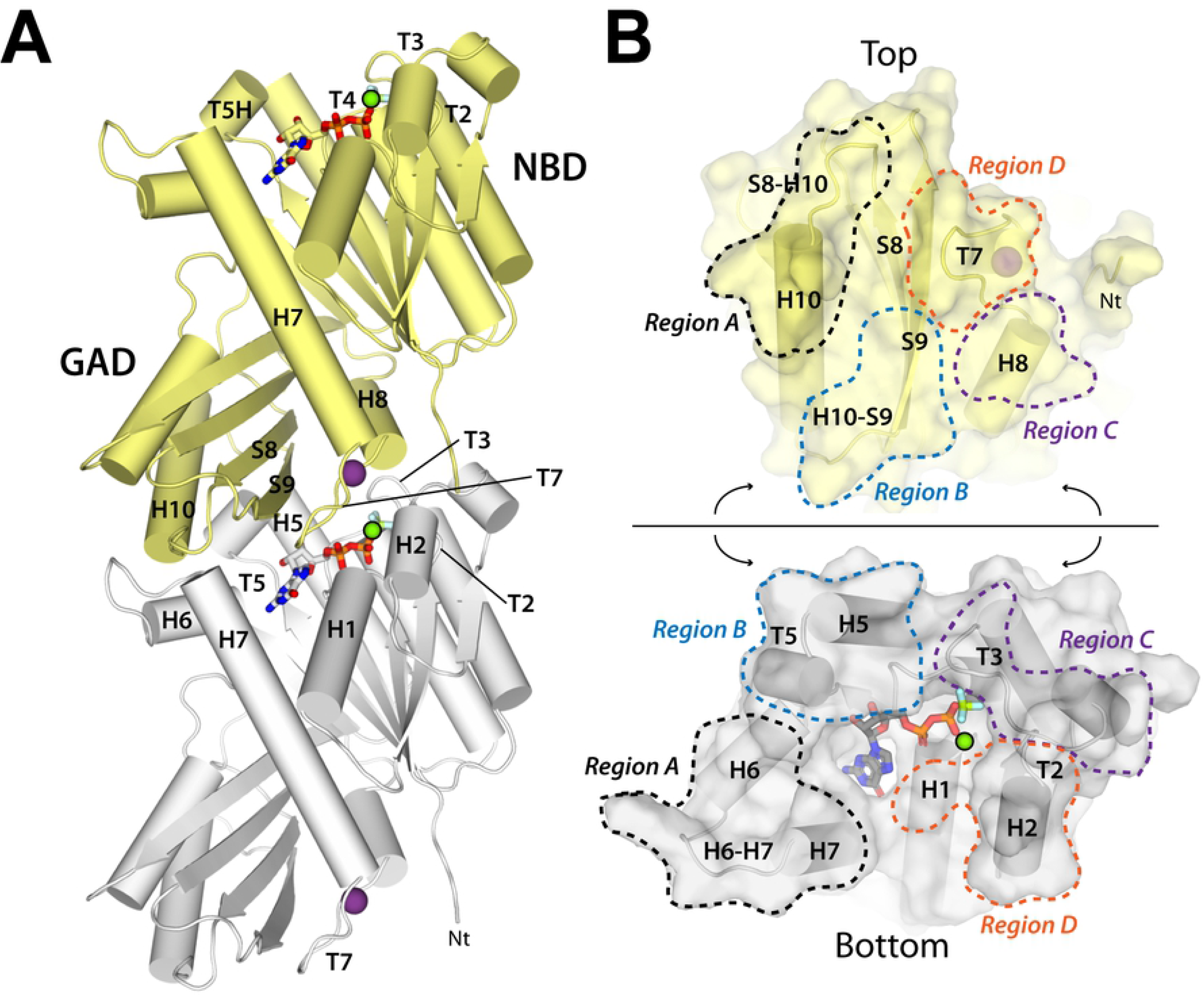
Filament structure of SaFtsZ complexed with GDP, BeF_3_^-^ and Mg^2+^. (A) Adjacent monomers, colored grey and yellow, in the FtsZ straight filaments are held by interactions around the nucleotide-binding pocket. The nucleotide-binding domain (NBD), the GTPase-activating domain (GAD) and relevant secondary structure elements are indicated. T5H is a mini-helix within loop T5, while monovalent and divalent ions appear as purple and green spheres. (B) Open view of the interface formed by adjacent monomers, showing relevant structural elements. Four major contact regions are marked with the same colour in each monomer.

### Structures mimicking the pre-catalytic state of GTP hydrolysis

The filament structure of SaFtsZ complexed to GDP and BeF_3_^-^ in the presence of Mg^2+^ reveals that BeF_3_^-^ adopts a tetrahedral configuration with beryllium lying 2.7 Å apart from GDP (Fig 3). Two fluorine atoms contact the bottom FtsZ monomer through hydrogen bonds (H-bond) with residues A71 and A73 in loop T3, G108 in loop T4, and T109 in helix H4 (Fig 3A). The third fluorine atom coordinates Mg^2+^, together with the β-phosphate of GDP. In a canonical pre-catalytic state bound to GTP, the positive charge of Mg^2+^ is expected to deprive in electrons the link between the β- and γ-phosphates in GTP, thus providing the main activating charge. Mg^2+^ can be replaced by other divalent cations, as shown by the structure in complex with Mn^2+^ (S3 Fig), which occupies the same binding site (S4A Fig). Comparison with the reported structure of GTP-bound SaFtsZ in the R conformation [14] suggests that switching into the filament-prone T conformation involves rearrangement of GTP and Mg^2+^ within their binding pocket (S4B Fig).

**Fig 3.**
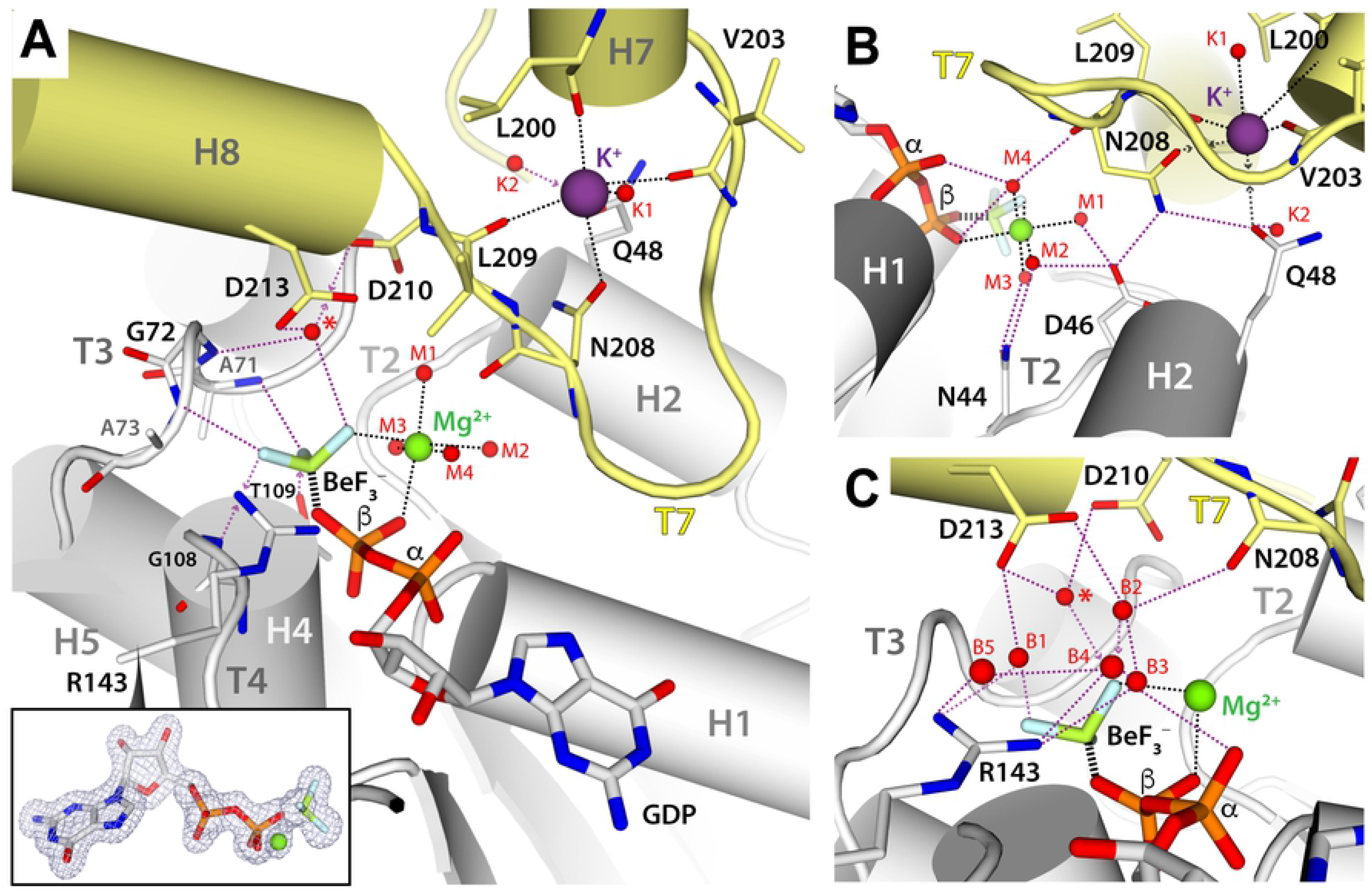
Nucleotide-binding pocket in the SaFtsZ structure complexed with GDP, BeF_3_^-^ and Mg^2+^. (A) Overall view of the interface between adjacent subunits. Inset: Polder OMIT map around the nucleotide mimetic and Mg^2+^ ion. (B) Zoom around the T7-H2 region. (C) Zoom around bridging water molecules. In all panels, structural elements and relevant residues are labelled, while Mg^2+^, K^+^ and water are represented as green, purple and red spheres, respectively. A red star indicates the pre-catalytic water. For clarity, certain solvent molecules, coordination contacts (black dash) and H-bonds (purple dash) are omitted in different panels.

An intricate network of solvent molecules surrounds BeF_3_^-^ and Mg^2+^ ions and contributes to stabilize the interactions between adjacent monomers. Four water molecules complete the octahedral coordination sphere of Mg^2+^ (Fig 3A). Three of these waters (M1-M3) form H-bonds with sidechains of N44 and D46 in loop T2 of the bottom monomer, while the fourth (M4) connects BeF_3_^-^ and the GDP α- and β-phosphates with the backbone of N208 in loop T7 of the top monomer (Fig 3B). In addition, BeF_3_^-^ establishes H-bonds with three other water molecules (Fig 3C): (i) the pre-catalytic water (Fig 3C, red star) held by residues D210 and D213 in the top T7 loop and by residue G72 in bottom loop T3; (ii) the water (B1) bridging sidechains of bottom residue R143 in helix H5 and top residue D213; and (iii) the water (B2) linking top residues D213 and N208. The last two solvent molecules belong to an H-bond connected path where three additional waters (B3-B5) bridge bottom residue R143 with the GDP *α*-phosphate and the backbone of top residues M292 and F294 (S2 Table). This solvent shell is altered in a structure lacking Mg^2+^, where bridging waters B2-B4 shift away from R143 by ∼1 Å and BeF_3_^-^ tilts away from the pre-catalytic water by 6°, likely weakening BeF_3_^-^ binding (S4C and S4D Fig).

While the N208 backbone connects to BeF_3_^-^ and Mg^2+^ ions via solvent molecules (M4 and B2), its sidechain both forms H-bonds with D46 and Q48 in the bottom monomer and coordinates a K^+^ ion within the top T7 loop (Fig 3B and 3C). The K^+^ ion is further coordinated by the backbone of residues L200, V203 and L209 in the top monomer, plus a solvent molecule (K1). Additionally, the sidechain of Q48 in bottom helix H2 lies 3.1 Å apart from K^+^, which is within the 3.6 Å limit of this cation [37], whereas a second water molecule (K2) connected to Q48 lies 4.2 Å away from the cation. The high resolution of our structures enabled us to assign density for this ion to K^+^ supported by results from various analysis, instead of a previously-assigned Ca^2+^ ion [25,29]. First, validation of atomic models with each of these ions using CheckMyMetal [38] favours K^+^. Second, crystals obtained in the presence of divalent cation chelators showed density for this ion that is identical to that observed in the original KCl buffer (S5 Fig). Third, free Ca^2+^ in solution is below 1 ion per 50 SaFtsZ molecules, as determined with a fluorescent probe (Materials and Methods). Finally, substitution of KCl by NaCl during protein purification yielded a structure where the T7 loop is occupied by Na^+^ (S5D Fig). The presence of Na^+^ in the T7 loop alters the configuration of ion-coordinating residues, especially V203. Notably, Q48 lies outside the Na^+^ coordination sphere whereas water K2 lies within and forms an H-bond with Q48. These changes, likely associated with increased filament stability, correlate with a tenfold reduction in GTPase activity upon substitution of KCl by NaCl (S3 Table).

We also sought to solve the structure of SaFtsZ complexed to GMPCPP, as it induces filament formation with Mg^2+^ (Fig 1C and 1D). However, the resulting structure only showed density for GMPCP, which binds SaFtsZ as GDP does [29] (S6A Fig), likely due to GMPCPP hydrolysis. In contrast, we determined the structure of SaFtsZ complexed to GMPPCP, where the C atom linking the β- and *γ*-phosphates in GMPPCP points in the opposite direction to that observed in the structure complexed to GDP and BeF_3_^-^ (S6B Fig). This configuration is incompatible with Mg^2+^ coordination, as confirmed by the structure of SaFtsZ complexed with GMPPCP and Mg^2+^ (S6C Fig). Unpredictably, however, Mg^2+^ enters a novel binding site located between the *γ*-phosphate and top residue D213. Binding of Mg^2+^ at this site, confirmed by solving the structure with Mn^2+^ (S3 Fig), alters the configuration of the interface contact between top T7 loop and bottom helix H2, such that Q48 moves away by 1 Å from loop T7 and K^+^ is evicted from its binding site (S6D Fig). This suggests that plasticity of the T7/H2 contact region (Region-D, Fig 2B) within straight filaments might influence cation exchange.

Overall, our results indicate that the GTP *γ*-phosphate provides a chemical environment for Mg^2+^ binding, reinforces interactions with the solvent network at the subunit interface, and enables bottom helix H2 to entrap a K^+^ cation within the top T7 loop.

### Structure mimicking the transition state of GTP hydrolysis

To gain further insight into the FtsZ catalytic mechanism, we determined the filament structure of SaFtsZ complexed to GDP, AlF_4_^-^ and Mg^2+^. The structure shows AlF_4_^-^ and Mg^2+^ in the canonical intermediate of catalysis (Fig 4A and S3 Fig), as described for classical GTPases [31,34]. AlF_4_^-^ adopts a planar configuration where the bond distance between aluminium and the GDP β-phosphate is 0.5 Å larger than that observed for BeF_3_^-^, with three fluorine atoms occupying equivalent positions and maintaining all protein and solvent interactions. The fourth fluorine atom in AlF_4_^-^ (Fig 4, grey star) points towards R143 and contacts waters B1-B2, which shift towards R143 by 0.6 and 0.8 Å respectively, while water B3 is absent. Notably, the catalytic water (Fig 4B, red star) approaches AlF_4_^-^ by 1.4 Å as compared to the BeF_3_^-^ structure, which is accompanied by a 30° rotation of catalytic residue D210. A squared bipyramid is, thus, formed between AlF_4_^-^, the linking oxygen in the GDP β-phosphate and the catalytic water. In addition, the geometry of K^+^ coordination is rearranged on both sides of the subunit interface. On one hand, the N208 sidechain contacts the Mg^2+^ coordination sphere while keeping direct contact with K^+^. On the other hand, the Q48 sidechain shifts 2.1 Å away from its K^+^ coordination position in the BeF_3_^-^ structure, while D46 moves 0.8 Å away from the Mg^2+^ ion. As a result, D46 loses an H-bond with the Mg^2+^ coordination sphere, compensated by H-bond formation with a new solvent molecule that also contacts top residue N208 (Fig 4C).

**Fig 4.**
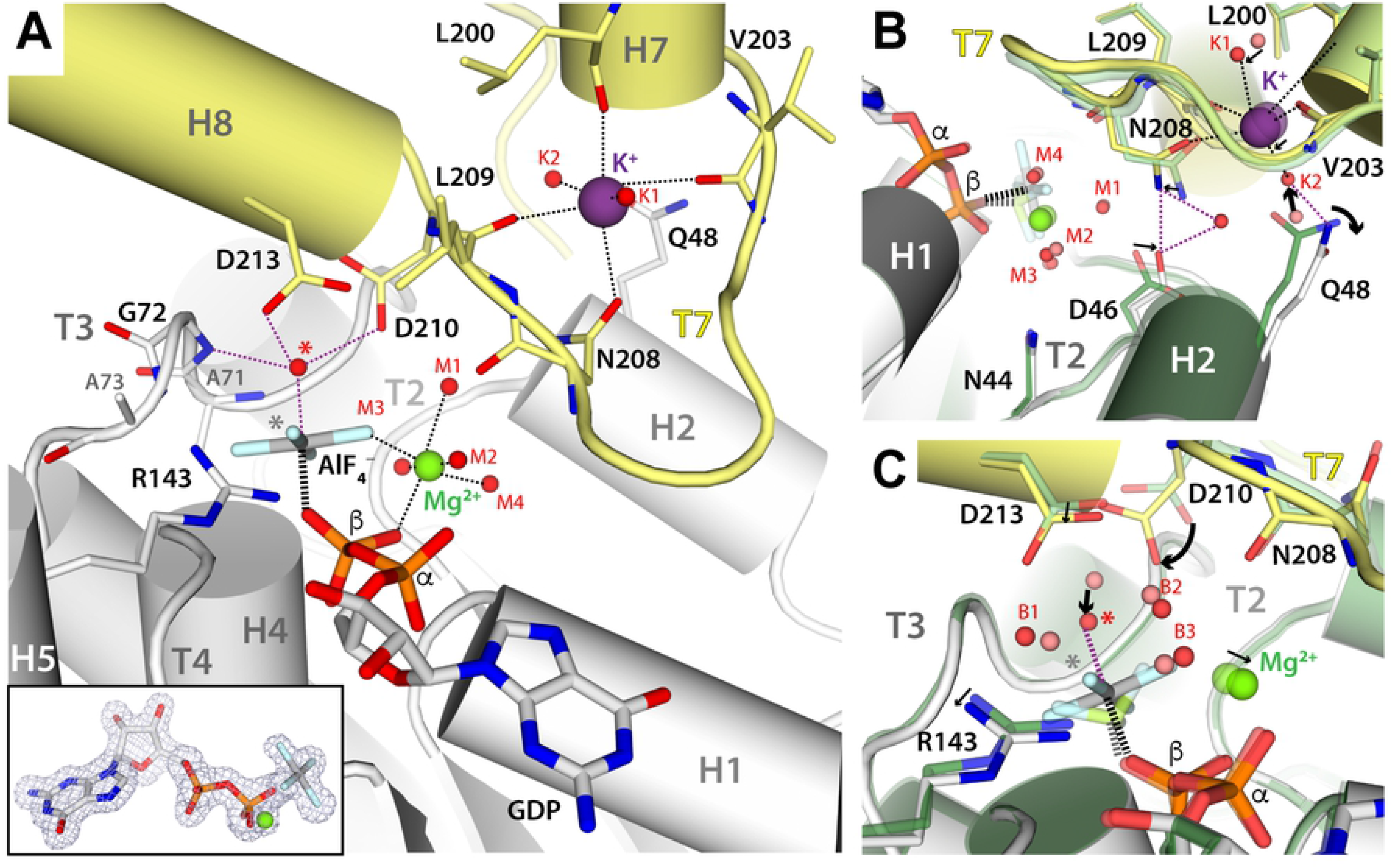
Nucleotide-binding pocket in the SaFtsZ structure complexed with GDP, AlF_4_^-^ and Mg^2+^. (A) Overall view of the interface between adjacent subunits. Structural elements and relevant residues are labelled, while Mg^2+^, K^+^ and water are represented as green, purple and red spheres, respectively. Red and grey stars indicate, respectively, the catalytic water and the fourth F atom position in AlF_4_^-^ as compared to BeF_3_^-^. Coordination contacts and H-bonds are shown as black and purple dash lines, respectively. Inset: Polder OMIT map around the nucleotide mimetic and Mg^2+^ ion. (B-C) Superposition of the SaFtsZ structure in complex with GDP, AlF_4_^-^ and Mg^2+^ (grey and yellow for bottom and top monomers, red spheres for water) onto that in complex with GDP, BeF_3_^-^ and Mg^2+^ (dark green and light green for bottom and top monomers, salmon spheres for water). Black arrows indicate conformational rearrangements between both structures.

### Nucleotide-dependent structural rearrangements and interfacial bonding

Differences between the BeF_3_^-^ and AlF_4_^-^ complexes at the residue level entail minor structural rearrangements at the subunit interface. The top T7 loop flattens towards the γ-phosphate position, while the neighbouring top GAD rotates by ∼5° towards top helix H7 (S1 and S2 Movies). This movement is coordinated with opening of bottom monomer motifs that surround the nucleotide, including helix H2, loop T3, loop T5, and the N-terminal end of helix H7. As a result, the interaction between top strand S9 and bottom loop T5 (Region-B, Fig 2B) is reinforced, while the contact between top loop T7 and bottom helix H2 (Region-D, Fig 2B) is weakened (S7A Fig).

Further comparison with the structure of SaFtsZ bound to only GDP in the same T conformation [29] shows that both the top and bottom monomers mostly recover the configuration observed in the BeF_3_^-^ structure (S3, S4, S5 and S6 Movies). Nevertheless, several interactions are absent in the GDP structure that likely destabilize the association interface. First, nine water-mediated H-bonds involving BeF_3_^-^ and Mg^2+^ ions are lost from the association interface, despite the presence of two solvent molecules in the BeF_3_^-^ site (S2 Table). Second, in contact Region-D (top T7/bottom H2), Q48 lies outside the K^+^ coordination sphere and the H-bond with top residue N208 is not restored (S7B and S7C Fig). Third, a salt bridge between bottom residue R67 and top residue D97 is missing in contact Region-C (S7D Fig). Besides, loss of an H-bond between GDP and bottom residue N25 in helix H1 likely weakens nucleotide binding by FtsZ (S7C Fig).

In all structures a pore, formed by the tip of the top T7 loop and residues from bottom helices H1 and H2, allows access into the BeF_3_^-^ site (S7E Fig). The pore, with a diameter of ∼2.5 Å and a length of ∼16 Å, allows Mg^2+^ exchange but is apparently too narrow for phosphate release. Nevertheless, BeF_3_^-^ could access the *γ*-phosphate location during crystal soaking (Materials and Methods), suggesting flexibility that would also allow phosphate release from the filament.

### Functional properties and structure of FtsZ interfacial mutants

Our structural studies identified D46, Q48, R143, N208, D210 and D213, all conserved across species, as key residues for subunit association or GTP hydrolysis. We constructed individual SaFtsZ mutants in these positions, which in EcFtsZ functionally inactivate the protein as observed using *in vivo* complementation tests [18]. Mutations D46A, R143Q, N208L, D210N and D213N, all suppress SaFtsZ assembly as monitored by light scattering and polymer sedimentation (Fig 5A and 5B), supporting a prominent role in filament formation. In accordance, the crystal structures of mutants D46A and D210N exhibit altered configurations at Region-D (Fig 5C). In the D46A mutant, bottom residue Q48 shifts 3 Å away from the K^+^ site, while the whole top T7 loop is distorted. In the D210N mutant, the top T7 loop lacks K^+^ and moves 2 Å away from the bottom monomer, while the bottom T3 loop is severely rearranged.

**Fig 5.**
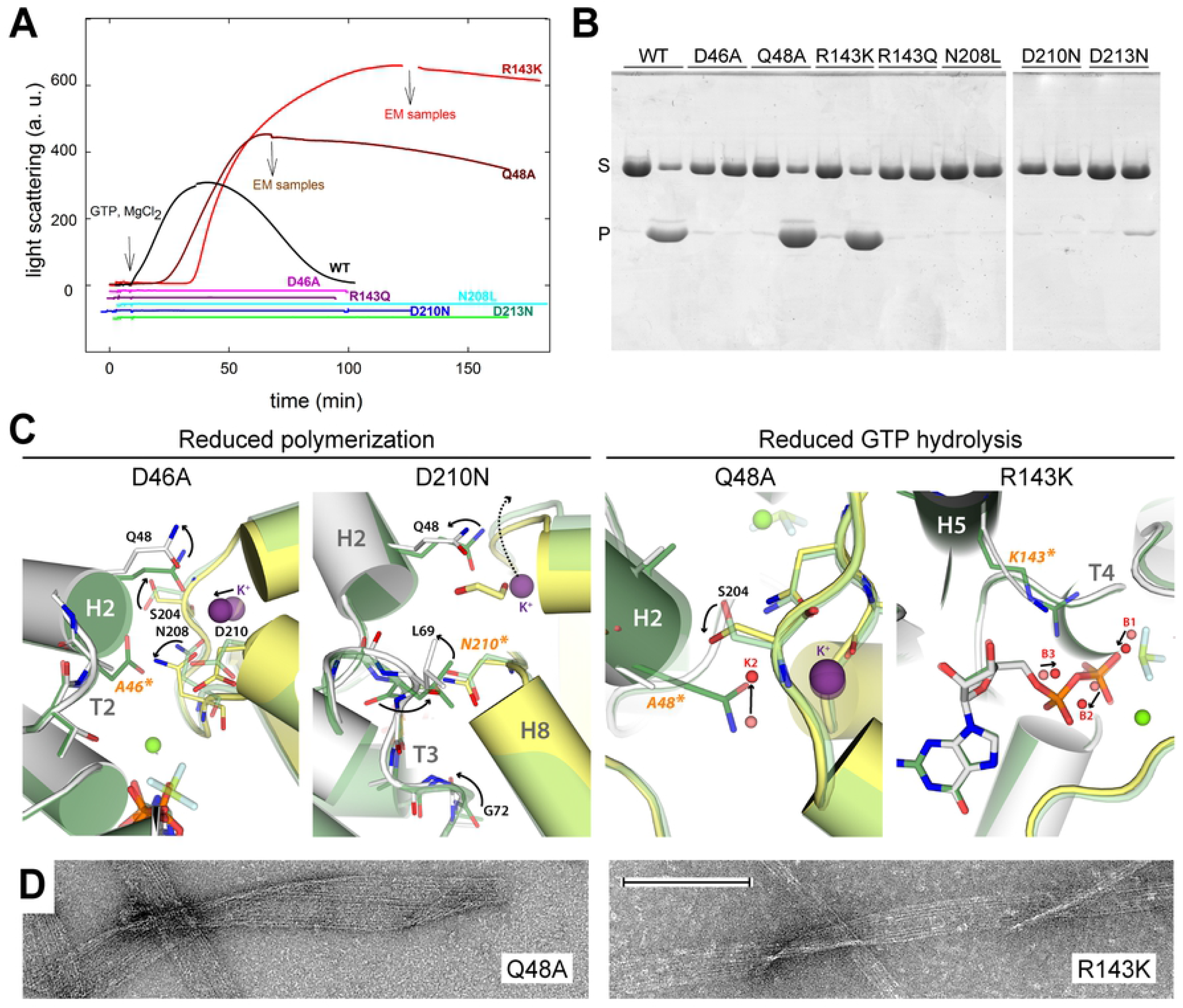
Functional and structural analysis of SaFtsZ mutants at the assembly interface. (A) Light scattering time courses during assembly of wild-type (black line), D46A, Q48A, R143Q, R143K, N208L, D210N and D213N (color lines) SaFtsZ (50 µM) to which 1 mM GTP was added and assembly triggered by addition of 10 mM MgCl_2_ at time zero. Experiments were made as in Fig. 1. (B) Sedimentation assays of polymer formation by wild-type and mutant SaFtsZ (31 µM) with 10 mM MgCl_2_ by high-speed centrifugation at the time of maximal scattering. For each sample, the left and right lanes contain 1 mM GDP and GTP, respectively; pellet (‘P’ label) and supernatant (‘S’ label) samples were consecutively loaded with a 15 min time shift in SDS-PAGE. (C) Superposition of SaFtsZ mutant structures in complex with GDP (grey and yellow for bottom and top monomers, red spheres for water) onto that of the wild-type protein in complex with GDP, BeF_3_^-^ and Mg^2+^ (dark green and light green for bottom and top monomers, salmon spheres for water). Black arrows indicate conformational rearrangements between both structures. (D) Representative electron micrographs of polymers formed by Q48A and R143K. The bar indicates 200 nm.

Interestingly, mutants Q48A and R143K supported assembly, forming polymers that are stabilized against disassembly by GTP consumption as compared to the wild-type protein (Fig 5A and 5B). This correlates with reduced GTPase activity by about six-fold in both mutants (S3 Table). The structure of the Q48A mutant shows that its K^+^ coordination position is occupied by water K2 (Fig 5C), indicating a role for this residue in securing K^+^ within the T7 loop for efficient catalysis. In the structure of the R143K mutant, K143 occupies an equivalent position to that of R143 in the wild-type protein but its positive charge lies 3 Å further from the GDP *α*-phosphate (Fig 5C). As the R143Q mutant is incompetent for polymerization and catalysis, our results indicate that a positively-charged residue in this position is required for GTP hydrolysis. The polymers formed by the Q48A and R143K mutants were mostly filamentous ribbons (Fig 5D), rather than the bundles observed for the wild-type protein (Fig 1D). Because the employed concentration of 8 mM Mg^2+^ slows polymer nucleation (notice lag times in Fig 5A), the effects of the mutations on assembly were confirmed at 2 mM Mg^2+^ (S8 Fig). While the wild-type protein forms rapidly-depolymerizing, single and double, wavy filaments, Q48A and R143K form stabilized, straight, thin filaments that slowly evolve into filamentous ribbons. These properties, which are strikingly similar to the effects of Na^+^ on wild type SaFtsZ assembly (S5 Fig), suggest that inhibition of GTP hydrolysis favors more rigid, straight SaFtsZ filaments that associate into ribbons.

## Discussion

Cytomotive proteins such as actins and tubulins self-assemble into nucleotide-hydrolyzing dynamic filaments that are able to perform mechanical work by themselves and also serve as rails for motor proteins [39]. Treadmilling of FtsZ filaments, driven by GTP hydrolysis, is essential for correct cell division in bacteria, which lack homologs of cytoskeletal motor proteins. In this work, we reveal the detailed mechanism of interfacial nucleotide hydrolysis by FtsZ filaments, by determining crystal filament structures in complex with different GTP mimetics and metal ions, combined with functional and structural characterization of individual mutants. Our results, together with previous structures, uncover the structural mechanism of FtsZ assembly dynamics at an unmatched resolution among cytomotive filaments.

### A composite active site with two labile metal ions

Our structures reveal new features of FtsZ as an atypical GTPase [20]. First, we identified swinging of D210 in the top T7 loop as a key event to position the catalytic water for attack over the *γ*-phosphate of GTP. In contrast, top residue D213 and bottom residue G72 essentially preserve their position during catalysis and rather provide a chemical environment for nucleophilic attack. Second, we found that bottom residue R143, which connects to D213 and the nucleotide *γ*-phosphate through bridging water B1, contributes to catalysis and that the position of its positive charge is critical for efficient GTP hydrolysis. Accordingly, R143K and not R143Q mutation allows assembly but presents reduced GTPase activity (Fig 5 and S3 Table). The equivalent residue in MjFtsZ, R169, was suggested to stabilize the transition state [12]. The role of R143 is likely equivalent to that of the arginine finger in Ras-like GTPases, while in these enzymes the finger residue contacts the nucleotide directly [40]. Third, we showed that a two-cation mechanism involving Mg^2+^ and K^+^ operates in FtsZ for GTP hydrolysis. While Mg^2+^ contacts the β- and *γ*-phosphates of GTP and has a direct role in catalysis, K^+^ locates in the top T7 loop and contacts the substrate indirectly through its coordinating residue N208, which forms H-bonds with solvent molecules connected to Mg^2+^ and the *γ*-phosphate. Accordingly, mutation N208L abolishes polymerization. Taken together, SaFtsZ can be classified as a type II Mg^2+^/K^+^ enzyme, as the monovalent cation has an allosteric role in catalysis [41]. This differs from most type I GTPases, where K^+^ assists catalysis through direct contact with the nucleotide phosphates, thus providing the activating charge that in other GTPases is supplied by the arginine finger [40]. Fourth, we found that locking of K^+^ within the top T7 loop by bottom residue Q48 plays a role in catalysis. In agreement, the Q48A mutant allows assembly but exhibits reduced GTPase activity (Fig 5 and S3 Table). K^+^ is bound weakly and can be evicted from its binding site by subtle changes around the T7 loop, as observed in the structures of mutant Q48A and wild-type SaFtsZ complexed to GMPPCP and Mg^2+^. Moreover, substitution of K^+^ by Na^+^ strongly reduces the GTPase activity (S5 Fig and S3 Table), similarly to FtsZ from other species, with some exceptions [42]. This K^+^ preference for hydrolysis correlates with estimated concentrations in the bacterial cytosol of 200 mM K^+^ and 5 mM Na^+^ [43].

### A dynamic water shell mediates assembly and catalysis

The high resolution of the structures reported here enables location of solvent molecules at the subunit interface that contribute to filament assembly and catalysis (S2 Table). Water molecules M1-M4, located between the nucleotide phosphates and the bottom T2 loop, coordinate Mg^2+^ binding, with water M4 playing a prominent role as it also bridges the nucleotide phosphates with the residue N208 in the top T7 loop (Fig 3). Notably, waters M1-M4 rearrange to accommodate the planar transition intermediate analogue AlF_4_^-^ and the sidechain of catalytic residue D210, thus enabling correct positioning of the catalytic water and minor opening of bottom residue Q48 and helix H2 (Fig 5B and S2 Movie). Besides, waters B1-B5 connect the nucleotide phosphates with the top T7 loop and strand S9. Moreover, relocation of waters B2-B4 upon Mg^2+^ binding enables bridging of R143 with the nucleotide phosphates, which arises as a key event to position its positive charge to assist catalysis. Furthermore, bridging waters B1-B3 rearrange to accommodate the catalysis intermediate, thus inducing slightly retraction of the R143 charge in the transition state. Solvent rearrangements also occur at the interface between bottom T2 and top T7 loops, which in the transition intermediate allow disengagement of bottom helix H2 from the top T7 loop.

### A crosstalk region modulates catalysis and disassembly

In spite of the rigid context of our structures, concerted rearrangements are observed between the structures complexed to BeF_3_^-^ or AlF_4_^-^ (S1 and S2 Movies), while additional changes arise in the absence of the *γ*-phosphate (S3, S4, S5 and S6 Movies). We distinguish two major interacting areas in the straight filament interface. On one hand, a pivoting area for interface opening, as observed in molecular dynamics simulations of SaFtsZ filaments [44], involves interface contact Regions A and B (Fig 2B). While rearranged, this area is roughly maintained in R conformation structures of SaFtsZ pseudofilaments [13,23,24]. On the other hand, we define a crosstalk area around the *γ*-phosphate site, which encompasses interface contact Regions C and D (Fig 2B) and is specific of straight filaments in the T conformation. The relevance of this crosstalk area is highlighted by mutational analysis showing that D46A, N208L, D210N, and D213N, all impair filament formation in solution. In agreement, the structures of mutants D46A and D210N exhibit an altered configuration around this area. A similar effect can be expected for N208 and D213N, which are both critical to maintain the T7 loop conformation and its interaction with the bottom monomer. Our structures in complex with GMPPCP in the presence and absence of Mg^2+^ further underscore the plasticity of the crosstalk interface within straight filaments.

### A model for FtsZ filament dynamics

We propose a model for the FtsZ catalytic assembly cycle (Fig 6). Our description starts with a filament where all subunits are in the T conformation and contain GTP. We reason that Mg^2+^ requirement for FtsZ polymerization is mainly related to its shielding effect over the acidic charge of the triphosphate nucleotide, while it also contributes to accurate positioning of the γ-phosphate for catalysis (Fig 3 and S4D Fig). Additionally, K^+^ within the filament T7 loop is secured through labile coordination with residue Q48 from the neighbouring subunit. In such configuration, nucleophilic attack by the catalytic water over the γ-phosphate occurs through an transition state where key residues and solvent molecules around the crosstalk region are rearranged. GTP hydrolysis proceeds at slow rate due to unique catalytic properties of FtsZ described above, followed by fast Mg^2+^ release through the exit pore, which likely rearranges to also liberate inorganic phosphate. This generates the assembled GDP-bound filament in the T conformation, where absence of key interfacial ionic interactions mediated by the γ-phosphate and Mg^2+^ destabilizes filament contacts, which is accompanied by minor structural rearrangements. We speculate that this allows detachment around the crosstalk region while contacts around the pivoting area are roughly maintained. This eventually allows structural relaxation into the R conformation, likely accompanied by K^+^ release from loop T7 as density for ion is absent across FtsZ structures in the R conformation, altogether leading to interfacial dissociation at the filament minus end. Free FtsZ monomers in the R conformation spontaneously exchange-in GTP and Mg^2+^ to enter a new assembly cycle at the filament plus end. The nucleotide *γ*-phosphate and Mg^2+^ likely stabilize helix H2 and loop T2 in the NBD of the R monomer such that its top surface is preorganised similar to the T conformation. However, the bottom surface of this R monomer is substantially different from the T conformation, including a highly-flexible T7 loop, whereas the exposed T7 loop in an all-T filament [15,16] likely adopts its association-prone configuration. Altogether, this suggests that the incoming R monomer should preferentially associate through its preorganised GTP-bound top surface with the bottom of the T filament exposing the T7 loop, which constitutes the growing end in agreement with reported mutational analysis of interface residues [17,18].

**Fig 6.**
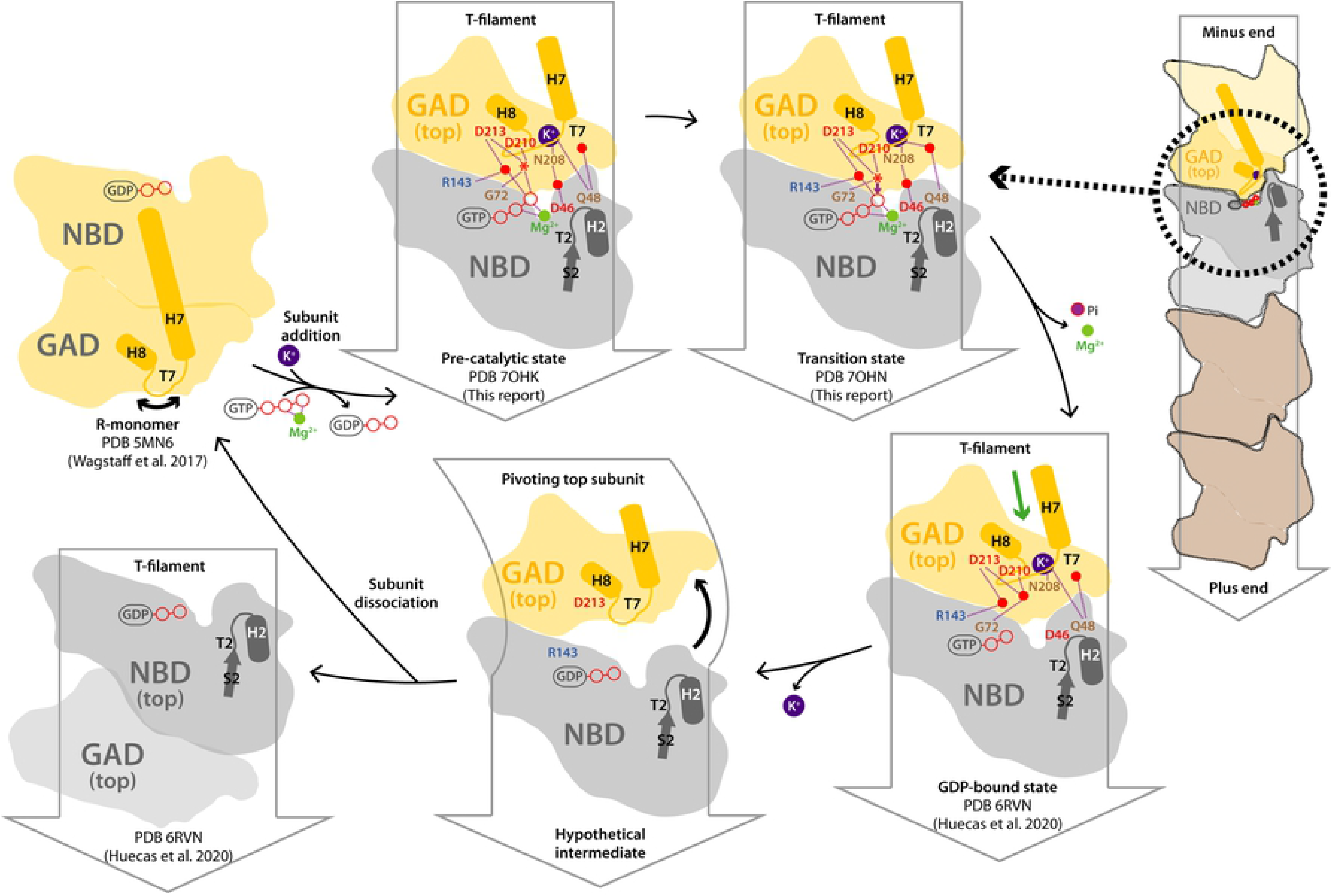
Model for FtsZ filament dynamics and catalytic assembly cycle. On the right, schematic FtsZ filament with top and second-top subunits in yellow and grey, respectively. The dotted black circle defines the area represented in the remaining cartoons of the filament assembly interface, where only the GAD of the top subunit and the NBD of second-top subunit are represented. Structural elements and relevant residues are labelled, while Mg^2+^, K^+^ and water are represented as green, purple and red spheres, respectively. A red star indicates the catalytic water, while H-bonds are shown as purple dashed lines. A green arrow indicates the cleft for PC190723 binding.

Our proposed mechanism unifies slow nucleotide hydrolysis with conformational switch and treadmilling, all essential properties for FtsZ function in bacterial cell division. It should be noticed that, analogously to FtsZ, tubulin [45] and actin [46,47] both show relatively small nucleotide-induced rearrangements associated to nucleotide hydrolysis in cryo-EM polymer structures, compared to the larger structural changes between their unassembled and assembled states. The primitive assembly mechanism of FtsZ might thus help to better understand those of the more complex cytomotive filaments.

## Materials and Methods

### Protein purification

The enzymatic core (residues 12-316) and full-length forms of FtsZ from methycillin-resistant *S. aureus* Mu50/ATCC 700699 (SaFtsZ and SaFtsZ_f_, respectively) complexed to GDP, as well as the nucleotide-free form (apoSaFtsZ), were all affinity-purified as described [29]. SaFtsZ single-residue mutants were prepared using QuikChange II Site-Directed Mutagenesis Kit (Agilent) with primers listed in S4 Table, and purified following the same protocol. For wild-type and mutant SaFtsZ, the final gel filtration was performed in 10 mM Tris-HCl pH 7.5, 50 mM KCl, 100 μM GDP. To obtain SaFtsZ in Na^+^ solution, KCl in the gel filtration buffer was replaced by 50 mM NaCl. To obtain SaFtsZ in complex with GTP, following thrombin cleavage, the sample was adjusted to 2 mM GTP before anion exchange in the absence of MgCl_2_, and adjusted again to 2 mM GTP before gel filtration where GDP was replaced by 100 µM GTP. Similarly, to produce SaFtsZ in complex with GMPPCP, the sample was adjusted to 100 µM GMPPCP before anion exchange in the absence of MgCl_2_, and adjusted again to 2 mM GMPPCP before gel filtration where GDP was replaced by 100 µM GMPPCP. To obtain SaFtsZ deprived of divalent cations in the solution, 1 mM EDTA was added to the gel filtration buffer. Full-length FtsZ from *E. coli* (EcFtsZ) was purified as described [48] and nucleotide-devoid, full-length FtsZ from *M. jannnaschii* (apo-MjFtsZ) was prepared as reported [49].

### Nucleotides, analogues and mimetics

GPD (sodium salt) and GTP (lithium salt) were from Sigma-Aldrich. GMPCP (Guanosine-5’-[(α,β)-methyleno]diphosphate, sodium salt), GMPCPP (Guanosine-5’-[(α,β)-methyleno]triphosphate, sodium salt) and GMPPCP (Guanosine-5’-[(β,γ)-methyleno]triphosphate, sodium salt) were from Jena Bioscience. Stock solutions of 80 mM BeF_3_^-^ were prepared by 2:3 mixing of BeSO_4_ with KHF_2_ neutralized with KOH while avoiding precipitation. The BeF_3_^-^ concentrations were calculated using the complex formation constants [50]. An AlF_3_/AlF_4_^-^ (1 mM) mixture was formed in by adding 1 mM AlCl_3_ and 2 mM KHF_2_ to protein or co-crystallization solutions. Predominantly AlF_4_^-^ (10 mM) was formed by adding 10 mM AlCl_3_ and 20 mM KHF_2_ [51], to crystal soaking solutions. The buffer pH was carefully adjusted as necessary.

### FtsZ assembly and GTPase activity

All FtsZ assembly experiments were made in 50 mM MES-KOH pH 6.5, 50 mM KCl, 1 mM EDTA (MES assembly buffer) at 25 °C, to which 10 mM MgCl_2_ and different nucleotide analogues were added, unless otherwise indicated. Polymer formation was monitored by right angle light scattering at 350 nm and by 30 min high-speed centrifugation followed by SDS-PAGE of pellet and supernatant samples and Coomassie blue staining. FtsZ polymers were visualized by negative stain electron microscopy, as described [29].

GTPase rate measurements were made from polymerizing SaFtsZ solutions (1 mM GTP, 10 mM MgCl_2_) by taking 10 µL aliquots every 3 min for the wild-type and every 15-30 min for the mutants, and measuring the released phosphate with the malachite green assay at 650 nm [52]. The hydrolysis rate values given are referred to total SaFtsZ concentration (50 µM).

### Measurement of Ca^2+^ concentration

Free Ca^2+^ concentrations were determined with the fluorescent probe Fura-FF pentapotassium salt (AAT Bioquest, Sunnnyvale, California). A Horiba Fluoromax-4 spectroflurometer was employed to measure the excitation spectra of probe (1 µM), with a 1 nm bandwith and emission set at 505 nm with 2 nm bandwith. A maximum at 360 nm was observed in the absence of Ca^2+^ that gradually shifted to a higher intensity maximum at 336 nm upon addition of increasing Ca^2+^ concentrations (1 to 100 µM) in the buffer. Comparison with these spectra showed that the free Ca^2+^ concentration was below 1 Ca^2+^ per 50 SaFtsZ molecules in the purified protein, < 1 µM in assembly (∼ 50 µM FtsZ) and crystallization (∼ 500 µM FtsZ) buffers and < 10 µM in crystallization solutions.

### Crystallization

Crystallization assays were carried out using purified SaFtsZ in 10 mM Tris-HCl pH 7.5, 300 mM KCl, 4% (v/v) 1-methyl-2-pyrrolidone, as described [29]. Crystals were grown at 295 K by vapor diffusion (sitting drop) under described conditions [25,29], i.e. 0.2 M lithium sulphate (Sigma), 10% (v/v) ethylene glycol (Merck, Darmstadt, Germany, for analysis), 0.1 M Tris-HCl pH 8.4–9.0, and 24–28% polyethylene glycol 5000 monomethyl ether (PEG5000; Aldrich, Steinheim, Germany).

The structure of SaFtsZ complexed with GDP and BeF_3_^-^ was obtained from crystals of 14 mg/ml SaFtsZ-GDP (28% PEG 5000 MME, pH 8.8) soaked in 10 mM BeF_3_^-^ during 20 h. The structure of SaFtsZ complexed to GDP, BeF_3_^-^ and Mg^2+^ was obtained by SaFtsZ-GDP co-crystallization with 2 mM BeF_3_^-^ and 10 mM MgCl_2_ (25% PEG 5000, pH 8.6). The structure of SaFtsZ complexed to GDP, BeF_3_^-^ and Mn^2+^ was obtained from SaFtsZ-GDP crystals (25% PEG 5000, pH 8.8) soaked in 10 mM BeF_3_^-^ and 20 mM MnCl_2_ during 20 h. The structure of SaFtsZ complexed to GDP and Na^+^ was obtained using SaFtsZ-GDP (15 mg/mL) purified with Na^+^, in 300 mM NaCl protein crystallization buffer (26% PEG 5000, pH 8.8). The structures of SaFtsZ-GDP in the presence of divalent cation chelators were obtained by soaking of SaFtsZ-GDP crystals in 10 mM CyDTA or 10 mM EGTA during 23 h, and by SaFtsZ-GDP crystallization in the presence of 1 mM EDTA (26% P(EG 5000, pH 8.5). The structure of SaFtsZ complexed to GDP, AlF_4_^-^ and Mg^2+^ was obtained from SaFtsZ-GDP crystals (24% PEG 5000, pH 8.6) soaked in 10 mM AlF_4_^-^ and 20 mM MgCl_2_ during 23 h. The structure of SaFtsZ complexed to GMPPCP was obtained by crystallization of 14 mg/ml SaFtsZ-GMPPCP-EDTA, with 1 mM GMPPCP, 190 μM EDTA and 216 μM MgCl_2_ (22% PEG 5000, pH 8.7). The FtsZ-GMPPCP-Mg^2+^ structure was obtained by soaking FtsZ-GMPPCP-EDTA crystals (22% PEG 5000, pH 8.8) in 20 mM MgCl_2_ during 23 h. The FtsZ-GMPPCP-Mn^2+^ structure was obtained by soaking FtsZ-GMPPCP-EDTA crystals (22% PEG 5000, pH 8.5) in 20 mM MnCl_2_ during 23 h. The structure of SaFtsZ complexed with GMPCP was obtained by slow co-crystallization of 25.5 mg/mL apo-SaFtsZ in 240 mM KAcetate (instead of 300 mM KCl) with 0.8 mM GMPCPP (26% PEG 5000, pH 8.8). The structure of SaFtsZ(D210N)-GDP was obtained from small crystals grown from SaFtsZ(210N)-GTP (15.6 mg/mL) in 27% PEG 5000, pH 8.8. The structure of SaFtsZ(R143K)-GDP (12.6 mg/mL) was from crystals grown in 24% PEG 5000, pH 8.4. The structure of SaFtsZ(Q48A)-GDP (15 mg/mL) was from crystals grown in 26% PEG 5000, pH 8.7. The structure of SaFtsZ(D46A)-GDP (8.7 mg/mL) was from crystals grown in 23% PEG 5000, pH 8.4.

### Structure determination

All crystals were flash-cooled by immersion in liquid nitrogen and diffraction data were collected at the ALBA (Spain) and ERSF (France) synchrotrons. All data were processed using XDS [53] and Aimless from the CCP4 Suite [54]. Data collection and refinement statistics are presented in S1 Table. The structures were determined through molecular replacement using Molrep [54] and the PDB entry 6RVN [29] as search model. Model building and refinement were done using Coot [55] and PHENIX [56], respectively. Refinement statistics are summarized in S1 Table. Structural figures were prepared using PyMOL (Schrödinger Inc.).

## Data availability

The structure factors and coordinates of all structures have been deposited in the Protein Data Bank (PDB) with the following accession codes: 7OHH (GDP+BeF_3_^-^), 7OHK (GDP+BeF_3_^-^ +Mg^2+^), 7OHL (GDP+BeF_3_^-^+Mn^2+^), 7OHN (GDP+AlF_4_^-^+Mg^2+^), 7OMJ (GMPPCP), 7OMP (GMPPCP+Mg^2+^), 7OMQ (GMPPCP+Mn^2+^), 7OJZ (GMPCP), 7OI2 (NaCl), 7ON2 (CyDTA), 7ON3 (EGTA), 7ON4 (EDTA), 7OJA (D210N mutant), 7OJB (R143K mutant), 7OJC (Q48A mutant), and 7OJD (D46A mutant).

## Acknowledgements

We thank Juan Estévez-Gallego and J. Fernando Díaz for sharing BeF_3_^-^ and AlF_x_ preparation protocols and discussions, David Juan for protein purification, and the Macromolecular Crystallography Facility at CIB. X-ray diffraction experiments were performed at ALBA and ESRF synchrotrons with help of local staff from beamlines XALOC and ID23-1, respectively.

## Supporting information captions

**S1 Fig. Assembly of full-length SaFtsZ is stabilized by BeF_3_^-^.** (A) Formation of polymers of full-length SaFtsZ (SaFtsZf, 49 µM), monitored by light scattering, with 1 mM GTP (blue line), with 1 mM GTP plus 5 mM BeF_3_^-^ (red line), with 1 mM GTP to which 1 mM BeF_3_^-^ was added at the point indicated by the arrow (green line), and with 1 mM GDP plus 5 mM BeF_3_^-^ (black line). In this experiment, 10 mM MgCl_2_ was added to the protein samples and assembly was triggered by nucleotide addition at time zero. The sample with GTP depolymerized upon nucleotide consumption (blue line), whereas the sample containing GTP plus 5 mM BeF_3_^-^ (red line) remained assembled, and disassembled with 20 mM EDTA (indicated by the red arrow). Addition of 5 mM BeF_3_^-^ to unassembled SaFtsZf with GDP (black line) or following disassembly by GTP hydrolysis (blue line and arrow) was not observed to induce polymerization nucleation in these solution experiments. (B) Representative electron micrograph (EM) of SaFtsZf polymers formed with GTP. (C) Polymers formed with GTP plus BeF_3_^-^. (D) Polymers formed with GMPCPP. (E) Small oligomers and protein aggregates with GMPPCP. EM samples were collected at maximum scattering in each case. Bar: 200 nm. Experiments were made in MES assembly buffer at 25 ⁰C. (F) Light scattering traces during assembly of full-length EcFtsZ (25 µM wild type protein) with 1mM GTP (blue line; 5 mM BeF_3_^-^ was added after depolymerization by nucleotide hydrolysis), with 1 mM GTP to which 5 mM BeF_3_^-^ was added at maximal light scattering (green line, later depolymerized by 20 mM EDTA), with 1 mM GTP plus 2 mM BeF_3_^-^ (pink line), with 1 mM GDP and 0.2 mM GTP plus 5 mM BeF_3_^-^ (red line), and with 1 mM GDP plus 5 mM BeF_3_^-^ (black line). (G) Light scattering assembly time course of full-length apo-MjFtsZ (15 µM) with 1mM GTP (blue line; 5 mM BeF3- added after depolymerization), with 1 mM GTP to which 5 mM BeF3- was added at maximal light scattering (green line), with 1 mM GDP plus 5 mM BeF3- and no GTP (black line), 10 µM GTP (brown line), 20 µM GTP (pink line), or 50 µM GTP (red line). Samples were depolymerized with 20 mM EDTA as indicated by blue and pink arrows. These assembly experiments with MjFtsZ were made at 55 ⁰C.

**S2 Fig. AlFx induces the formation of aberrant SaFtsZ polymers.** (A) Light scattering was employed to monitor the formation of polymers by the SaFtsZ core (50 µM) with 1 mM GTP (blue line), with 1 mM GTP plus 1 mM AlFx (red line), with 1 mM GDP plus 1 mM AlFx (magenta line) and with 1 mM GDP plus 0.1 mM AlFx (green line). Assembly was triggered by addition of 10 mM MgCl_2_ at time zero. Samples stabilized with 1 mM AlFx could be disassembled with 20 mM EDTA divalent metal chelator (indicated by the arrows). The effects of adding AlFx to polymers preassembled with GTP could not be determined due to precipitate formation. (B) Electron micrograph of polymers formed with 1 mM GTP. (C) Polymers formed with 1 mM GTP + 1 mM AlFx. (D) Enlarged view with 1 mM GTP + 1 mM AlFx. (E) Polymers formed with 1 mM GDP + 1 mM AlFx. (F) Polymers formed with 1 mM GDP + 0.1 mM AlFx. (G) Polymers formed with 1 mM GDP + 1 mM AlCl_3_ control. The bars indicate 200 nm.

**S3 Fig. Polder OMIT maps of SaFtsZ structures in complex with different GTP mimetics.** Expanded view of the nucleotide binding pocket, with polder OMIT electron density maps (blue mesh) contoured at 3 sigma around the different GTP mimetics and ions.

**S4 Fig. Comparison of SaFtsZ complexed with GDP, BeF_3_^-^ and Mg^2+^ to other structures.** In all panels, the SaFtsZ complexed with GDP, BeF_3_^-^ and Mg^2+^ is shown in grey and water molecules in this structure appear as red spheres. (A) Superposition onto the filament structure of SaFtsZ complexed with GDP, BeF_3_^-^ and Mn^2+^ (dark green for protein, salmon spheres for water). (B) Superposition onto the structure of SaFtsZ complexed with GTP and Mg^2+^ in the R conformation (PDB 5MN7, dark green). (C) Superposition onto the filament structure of SaFtsZ complexed with GDP and BeF_3_^-^ in the absence of Mg^2+^ (dark green for protein, salmon spheres for water). (D) Close-up view of panel C superposition around BeF_3_^-^ and the *β*-phosphate.

**S5 Fig. Confirmation of K^+^ binding at the T7 loop.** (A) SaFtsZ structure in complex with GDP, crystallized in the presence of 1 mM EDTA and 50 mM KCl. (B) SaFtsZ structure in complex with GDP of crystals soaked in 10 mM EGTA and 50 mM KCl. (C) SaFtsZ structure in complex with GDP of crystals soaked in 10 mM CyDTA and 50 mM KCl. (D) SaFtsZ structure in complex with GDP crystallized in the presence of 150 mM NaCl refined with Na^+^ (left and middle panels) or K^+^ (right panel) in the T7 loop. In all cases, 2Fo-Fc (blue) and Fo-Fc (green for positive, red for negative values) maps are contoured at 1.5 and 3.0 sigma, respectively. In the left panel, the structure (yellow and grey) is superposed to that in complex with GDP, BeF_3_^-^ and Mg^2+^. (E) Light scattering assembly time courses of SaFtsZ (50 µM) in K^+^ (black lines) and Na^+^-containing (pink lines) MES assembly buffers at 25 ⁰C. 1 mM GTP was added, and assembly was triggered by addition of 10 mM MgCl_2_ (solid lines) or 5 mM MgCl_2_ (dashed lines) as indicated by the arrow. (F) Representative electron micrograph of SaFtsZ in Na^+^ buffer with 10 mM MgCl_2_, to be compared with polymers formed in K^+^ buffer (Figure 1C). The bar indicates 200 nm. Notice the similarity of these scattering and electron microscopy results of SaFtsZ in Na^+^ buffer with those of the Q48A and R143K mutants in K^+^ buffer (Figures 6 and S8).

**S6 Fig. Structures of SaFtsZ complexed with GMPCP or GMPPCP.** Bottom and top monomers are gray and yellow, while dark and light green is used for bottom and top monomers of structures complexed to BeF_3_^-^. Water molecules are red, while salmon is used for solvent molecules in structures complexed to BeF_3_^-^. Mg^2+^ and K^+^ appear as green and purple spheres, respectively. (A) Structure complexed to GMPCP superimposed onto that complexed with GDP, both in the absence of Mg^2+^. (B) Structure complexed to GMPPCP superimposed onto that in complex with GDP and BeF_3_^-^, both in the absence of Mg^2+^. (C) Structure complexed to GMPPCP superimposed onto that in complex with GDP and BeF_3_^-^, both in the presence of Mg^2+^. (D) Close-up view of the structure complexed to GMPPCP and Mg^2+^. Coordination contacts and H-bonds are shown as black and purple dash lines, respectively.

**S7 Fig. Conformational rearrangements between different states of GTP hydrolysis.** (A) Superposition of the SaFtsZ structure in complex with GDP, BeF_3_^-^ and Mg^2+^ (grey and yellow for bottom and top monomers, red spheres for water) onto that in complex with GDP, AlF_4_^-^ and Mg^2+^ (green and light green for bottom and top monomers, salmon spheres for water). Black arrows indicate conformational rearrangements between both structures. (B) Superposition of the SaFtsZ structure in complex with GDP, AlF_4_^-^ and Mg^2+^ (green and light green for bottom and top monomers, salmon spheres for water) onto that in complex with GDP alone (purple and pink for bottom and top monomers, violet spheres for water). (C) Superposition of the SaFtsZ structure in complex with GDP, BeF_3_^-^ and Mg^2+^ (grey and yellow for bottom and top monomers, red spheres for water) onto that in complex with GDP alone (purple and pink for bottom and top monomers, violet spheres for water). (D) Superposition of the three structures in panels A-C around residue R67. (E) Nucleotide-binding pocket (pink) formed at the interface between adjacent monomers, showing a pore from the solvent into the Mg^2+^ binding site. T5H is a mini-helix within loop T5.

**S8 Fig. Assembly of SaFtsZ interfacial mutants in low Mg^2+^.** (A) Light scattering time courses during assembly of wild type protein (black trace, 50 µM), D46A, Q48A, R143Q, R143K, N201A, D210N and D213N (color traces, 50 µM each) in MES assembly buffer at 25 ⁰C, to which 2 mM GTP was added; assembly was triggered by addition of 5 mM MgCl_2_ at time zero. (B) Negatively stained electron micrograph of wild type SaFtsZ polymers, at maximum light scattering. (C) Mutant Q48A at the intermediate scattering plateau. (D) R143K at the intermediate scattering plateau. (E) Q48A at maximum scattering. (F) R143A at maximum scattering. The bar indicates 200 nm.

**S1 Table. Data collection and refinement statistics.**

**S2 Table. Water molecules at the subunit interface.**

**S3 Table. GTPase activity under assembly conditions.**

**S4 Table. Primers used in this study.**

**S1 Movie. Comparison between ground and transition states (overall).** Overall structural rearrangements at the subunit interface in the switch from BeF_3_^-^ (ground state mimic) to AlF_4_^-^ (transition state mimic) structures. The movement is repeated five times.

**S2 Movie. Comparison between ground and transition states (detail).** Detailed structural rearrangements around the active site in the switch from BeF_3_^-^ (ground state mimic) to AlF_4_^-^ (transition state mimic) structures.

**S3 Movie. Comparison between transition and phosphate-free states (overall).** Overall structural rearrangements at the subunit interface in the switch from AlF_4_^-^ (transition state mimic) to only-GDP (phosphate-free state) structures. The movement is repeated five times.

**S4 Movie. Comparison between transition and phosphate-free states (detail).** Detailed structural rearrangements around the active site in the switch from AlF_4_^-^ (transition state mimic) to only-GDP (phosphate-free state) structures.

**S5 Movie. Comparison between ground and phosphate-free states (overall).** Overall structural rearrangements at the subunit interface from BeF_3_^-^ (ground state mimic) to only-GDP (phosphate-free state) structures. The movement is repeated five times.

**S6 Movie. Comparison between ground and phosphate-free states (detail).** Detailed structural rearrangements around the active site from BeF_3_^-^ (ground state mimic) to only-GDP (phosphate-free state) structures. The BeF_3_^-^ and Mg^2+^ ions have been omitted for clarity.

## References

1. Bi EF, Lutkenhaus J. FtsZ ring structure associated with division in Escherichia coli. Nature. 1991;354: 161–164. doi:10.1038/354161a0

2. Stricker J, Maddox P, Salmon ED, Erickson HP. Rapid assembly dynamics of the Escherichia coli FtsZ-ring demonstrated by fluorescence recovery after photobleaching. Proc Natl Acad Sci U S A. 2002;99: 3171–3175. doi:10.1073/pnas.052595099

3. McQuillen R, Xiao J. Insights into the Structure, Function, and Dynamics of the Bacterial Cytokinetic FtsZ-Ring. Annu Rev Biophys. 2020;49: 309–341. doi:10.1146/annurev-biophys-121219-081703

4. Barrows JM, Goley ED. FtsZ dynamics in bacterial division: What, how, and why? Curr Opin Cell Biol. 2021;68: 163–172. doi:10.1016/j.ceb.2020.10.013

5. den Blaauwen T, Andreu JM, Monasterio O. Bacterial cell division proteins as antibiotic targets. Bioorg Chem. 2014;55: 27–38. doi:10.1016/j.bioorg.2014.03.007

6. Loose M, Mitchison TJ. The bacterial cell division proteins FtsA and FtsZ self-organize into dynamic cytoskeletal patterns. Nat Cell Biol. 2014;16: 38–46. doi:10.1038/ncb2885

7. Ramirez-Diaz DA, García-Soriano DA, Raso A, Mücksch J, Feingold M, Rivas G, et al. Treadmilling analysis reveals new insights into dynamic FtsZ ring architecture. PLoS Biol. 2018;16: e2004845. doi:10.1371/journal.pbio.2004845

8. Bisson-Filho AW, Hsu Y-P, Squyres GR, Kuru E, Wu F, Jukes C, et al. Treadmilling by FtsZ filaments drives peptidoglycan synthesis and bacterial cell division. Science. 2017;355: 739–743. doi:10.1126/science.aak9973

9. Yang X, Lyu Z, Miguel A, McQuillen R, Huang KC, Xiao J. GTPase activity-coupled treadmilling of the bacterial tubulin FtsZ organizes septal cell wall synthesis. Science. 2017;355: 744–747. doi:10.1126/science.aak9995

10. Monteiro JM, Pereira AR, Reichmann NT, Saraiva BM, Fernandes PB, Veiga H, et al. Peptidoglycan synthesis drives an FtsZ-treadmilling-independent step of cytokinesis. Nature. 2018;554: 528–532. doi:10.1038/nature25506

11. Scheffers D-J, de Wit JG, den Blaauwen T, Driessen AJM. GTP hydrolysis of cell division protein FtsZ: evidence that the active site is formed by the association of monomers. Biochemistry. 2002;41: 521–529. doi:10.1021/bi011370i

12. Oliva MA, Cordell SC, Löwe J. Structural insights into FtsZ protofilament formation. Nat Struct Mol Biol. 2004;11: 1243–1250. doi:10.1038/nsmb855

13. Fujita J, Harada R, Maeda Y, Saito Y, Mizohata E, Inoue T, et al. Identification of the key interactions in structural transition pathway of FtsZ from Staphylococcus aureus. J Struct Biol. 2017;198: 65–73. doi:10.1016/j.jsb.2017.04.008

14. Wagstaff JM, Tsim M, Oliva MA, García-Sanchez A, Kureisaite-Ciziene D, Andreu JM, et al. A Polymerization-Associated Structural Switch in FtsZ That Enables Treadmilling of Model Filaments. mBio. 2017;8. doi:10.1128/mBio.00254-17

15. Huecas S, Llorca O, Boskovic J, Martín-Benito J, Valpuesta JM, Andreu JM. Energetics and geometry of FtsZ polymers: nucleated self-assembly of single protofilaments. Biophys J. 2008;94: 1796–1806. doi:10.1529/biophysj.107.115493

16. Miraldi ER, Thomas PJ, Romberg L. Allosteric models for cooperative polymerization of linear polymers. Biophys J. 2008;95: 2470–2486. doi:10.1529/biophysj.107.126219

17. Du S, Pichoff S, Kruse K, Lutkenhaus J. FtsZ filaments have the opposite kinetic polarity of microtubules. Proc Natl Acad Sci U S A. 2018;115: 10768–10773. doi:10.1073/pnas.1811919115

18. Redick SD, Stricker J, Briscoe G, Erickson HP. Mutants of FtsZ targeting the protofilament interface: effects on cell division and GTPase activity. J Bacteriol. 2005;187: 2727–2736. doi:10.1128/JB.187.8.2727-2736.2005

19. Corbin LC, Erickson HP. A Unified Model for Treadmilling and Nucleation of Single-Stranded FtsZ Protofilaments. Biophys J. 2020;119: 792–805. doi:10.1016/j.bpj.2020.05.041

20. Nogales E, Downing KH, Amos LA, Löwe J. Tubulin and FtsZ form a distinct family of GTPases. Nat Struct Biol. 1998;5: 451–458. doi:10.1038/nsb0698-451

21. Löwe J, Amos LA. Crystal structure of the bacterial cell-division protein FtsZ. Nature. 1998;391: 203–206. doi:10.1038/34472

22. Oliva MA, Trambaiolo D, Löwe J. Structural insights into the conformational variability of FtsZ. J Mol Biol. 2007;373: 1229–1242. doi:10.1016/j.jmb.2007.08.056

23. Schumacher MA, Ohashi T, Corbin L, Erickson HP. High-resolution crystal structures of Escherichia coli FtsZ bound to GDP and GTP. Acta Crystallogr F Struct Biol Commun. 2020;76: 94–102. doi:10.1107/S2053230X20001132

24. Yoshizawa T, Fujita J, Terakado H, Ozawa M, Kuroda N, Tanaka SI, et al. Crystal structures of the cell-division protein FtsZ from Klebsiella pneumoniae and Escherichia coli. Acta Crystallogr F Struct Biol Commun. 2020;76: 86–93. doi:10.1107/S2053230X2000076X

25. Matsui T, Yamane J, Mogi N, Yamaguchi H, Takemoto H, Yao M, et al. Structural reorganization of the bacterial cell-division protein FtsZ from Staphylococcus aureus. Acta Crystallogr D Biol Crystallogr. 2012;68: 1175–1188. doi:10.1107/S0907444912022640

26. Tan CM, Therien AG, Lu J, Lee SH, Caron A, Gill CJ, et al. Restoring methicillin-resistant Staphylococcus aureus susceptibility to β-lactam antibiotics. Sci Transl Med. 2012;4: 126ra35. doi:10.1126/scitranslmed.3003592

27. Huecas S, Araújo-Bazán L, Ruiz FM, Ruiz-Ávila LB, Martínez RF, Escobar-Peña A, et al. Targeting the FtsZ Allosteric Binding Site with a Novel Fluorescence Polarization Screen, Cytological and Structural Approaches for Antibacterial Discovery. J Med Chem. 2021;64: 5730–5745. doi:10.1021/acs.jmedchem.0c02207

28. Matsui T, Han X, Yu J, Yao M, Tanaka I. Structural change in FtsZ Induced by intermolecular interactions between bound GTP and the T7 loop. J Biol Chem. 2014;289: 3501–3509. doi:10.1074/jbc.M113.514901

29. Huecas S, Canosa-Valls AJ, Araújo-Bazán L, Ruiz FM, Laurents DV, Fernández-Tornero C, et al. Nucleotide-induced folding of cell division protein FtsZ from Staphylococcus aureus. FEBS J. 2020;287: 4048–4067. doi:10.1111/febs.15235

30. Bigay J, Deterre P, Pfister C, Chabre M. Fluoride complexes of aluminium or beryllium act on G-proteins as reversibly bound analogues of the gamma phosphate of GTP. EMBO J. 1987;6: 2907–2913.

31. Jin Y, Richards NG, Waltho JP, Blackburn GM. Metal Fluorides as Analogues for Studies on Phosphoryl Transfer Enzymes. Angew Chem Int Ed Engl. 2017;56: 4110–4128. doi:10.1002/anie.201606474

32. Carlier MF, Didry D, Melki R, Chabre M, Pantaloni D. Stabilization of microtubules by inorganic phosphate and its structural analogues, the fluoride complexes of aluminum and beryllium. Biochemistry. 1988;27: 3555–3559. doi:10.1021/bi00410a005

33. Estévez-Gallego J, Josa-Prado F, Ku S, Buey RM, Balaguer FA, Prota AE, et al. Structural model for differential cap maturation at growing microtubule ends. Elife. 2020;9: e50155. doi:10.7554/eLife.50155

34. Wittinghofer A. Signaling mechanistics: aluminum fluoride for molecule of the year. Curr Biol. 1997;7: R682–685. doi:10.1016/s0960-9822(06)00355-1

35. Mingorance J, Tadros M, Vicente M, González JM, Rivas G, Vélez M. Visualization of single Escherichia coli FtsZ filament dynamics with atomic force microscopy. J Biol Chem. 2005;280: 20909–20914. doi:10.1074/jbc.M503059200

36. Huecas S, Ramírez-Aportela E, Vergoñós A, Núñez-Ramírez R, Llorca O, Díaz JF, et al. Self-Organization of FtsZ Polymers in Solution Reveals Spacer Role of the Disordered C-Terminal Tail. Biophys J. 2017;113: 1831–1844. doi:10.1016/j.bpj.2017.08.046

37. Harding MM. Metal-ligand geometry relevant to proteins and in proteins: sodium and potassium. Acta Crystallogr D Biol Crystallogr. 2002;58: 872–874. doi:10.1107/s0907444902003712

38. Zheng H, Cooper DR, Porebski PJ, Shabalin IG, Handing KB, Minor W. CheckMyMetal: a macromolecular metal-binding validation tool. Acta Crystallogr D Struct Biol. 2017;73: 223–233. doi:10.1107/S2059798317001061

39. Aylett CHS, Löwe J, Amos LA. New insights into the mechanisms of cytomotive actin and tubulin filaments. Int Rev Cell Mol Biol. 2011;292: 1–71. doi:10.1016/B978-0-12-386033-0.00001-3

40. Shalaeva DN, Cherepanov DA, Galperin MY, Golovin AV, Mulkidjanian AY. Evolution of cation binding in the active sites of P-loop nucleoside triphosphatases in relation to the basic catalytic mechanism. Elife. 2018;7. doi:10.7554/eLife.37373

41. Di Cera E. A structural perspective on enzymes activated by monovalent cations. J Biol Chem. 2006;281: 1305–1308. doi:10.1074/jbc.R500023200

42. Lu C, Stricker J, Erickson HP. FtsZ from Escherichia coli, Azotobacter vinelandii, and Thermotoga maritima--quantitation, GTP hydrolysis, and assembly. Cell Motil Cytoskeleton. 1998;40: 71–86. doi:10.1002/(SICI)1097-0169(1998)40:1<71::AID-CM7>3.0.CO;2-I

43. Shabala L, Bowman J, Brown J, Ross T, McMeekin T, Shabala S. Ion transport and osmotic adjustment in Escherichia coli in response to ionic and non-ionic osmotica. Environ Microbiol. 2009;11: 137–148. doi:10.1111/j.1462-2920.2008.01748.x

44. Ramírez-Aportela E, López-Blanco JR, Andreu JM, Chacón P. Understanding nucleotide-regulated FtsZ filament dynamics and the monomer assembly switch with large-scale atomistic simulations. Biophys J. 2014;107: 2164–2176. doi:10.1016/j.bpj.2014.09.033

45. Zhang R, LaFrance B, Nogales E. Separating the effects of nucleotide and EB binding on microtubule structure. Proc Natl Acad Sci U S A. 2018;115: E6191–E6200. doi:10.1073/pnas.1802637115

46. Merino F, Pospich S, Funk J, Wagner T, Küllmer F, Arndt H-D, et al. Structural transitions of F-actin upon ATP hydrolysis at near-atomic resolution revealed by cryo-EM. Nat Struct Mol Biol. 2018;25: 528–537. doi:10.1038/s41594-018-0074-0

47. Chou SZ, Pollard TD. Mechanism of actin polymerization revealed by cryo-EM structures of actin filaments with three different bound nucleotides. Proc Natl Acad Sci U S A. 2019;116: 4265–4274. doi:10.1073/pnas.1807028115

48. Rivas G, López A, Mingorance J, Ferrándiz MJ, Zorrilla S, Minton AP, et al. Magnesium-induced linear self-association of the FtsZ bacterial cell division protein monomer. The primary steps for FtsZ assembly. J Biol Chem. 2000;275: 11740–11749. doi:10.1074/jbc.275.16.11740

49. Huecas S, Schaffner-Barbero C, García W, Yébenes H, Palacios JM, Díaz JF, et al. The interactions of cell division protein FtsZ with guanine nucleotides. J Biol Chem. 2007;282: 37515–37528. doi:10.1074/jbc.M706399200

50. Mesmer RE, Baes CF. Fluoride complexes of beryllium(II) in aqueous media. Inorg Chem. 1969;8: 618–626. doi:10.1021/ic50073a042

51. Goldstein Gerald. Equilibrium Distribution of Metal-Fluoride Complexes. Anal Chem. 1964;36: 243–244. doi:10.1021/ac60207a074

52. Kodama T, Fukui K, Kometani K. The initial phosphate burst in ATP hydrolysis by myosin and subfragment-1 as studied by a modified malachite green method for determination of inorganic phosphate. J Biochem. 1986;99: 1465–1472. doi:10.1093/oxfordjournals.jbchem.a135616

53. Kabsch W. XDS. Acta Crystallogr D Biol Crystallogr. 2010;66: 125–132. doi:10.1107/S0907444909047337

54. Winn MD, Ballard CC, Cowtan KD, Dodson EJ, Emsley P, Evans PR, et al. Overview of the CCP4 suite and current developments. Acta Crystallogr D Biol Crystallogr. 2011;67: 235–242. doi:10.1107/S0907444910045749

55. Emsley P, Lohkamp B, Scott WG, Cowtan K. Features and development of Coot. Acta Crystallogr D Biol Crystallogr. 2010;66: 486–501. doi:10.1107/S0907444910007493

56. Adams PD, Afonine PV, Bunkóczi G, Chen VB, Davis IW, Echols N, et al. PHENIX: a comprehensive Python-based system for macromolecular structure solution. Acta Crystallogr D Biol Crystallogr. 2010;66: 213–221. doi:10.1107/S0907444909052925

